# A Meier-Gorlin Syndrome Mutation in Orc4 Causes Tissue-Specific DNA Replication Defects in *Drosophila melanogaster*

**DOI:** 10.1101/711820

**Authors:** Stephen L. McDaniel, Anna M. Branstad, Allison J. Hollatz, Catherine A. Fox, Melissa M. Harrison

**Affiliations:** University of Wisconsin School of Medicine and Public Health, Department of Biomolecular Chemistry, Madison, WI 53706

**Keywords:** *Drosophila*, Meier-Gorlin syndrome, Disease Model, DNA Replication

## Abstract

Meier-Gorlin syndrome is a rare recessive disorder characterized by a number of distinct developmental defects, including primordial dwarfism, small ears, and small or missing patella. Genes encoding members of the origin recognition complex (ORC) and additional proteins essential for DNA replication (CDC6, CDT1, GMNN, CDC45, and MCM5) are mutated in individuals diagnosed with MGS. The primary role of ORC is to license origins during the G1 phase of the cell cycle, but it also plays roles in cilia development, heterochromatin formation, and other cellular processes. Because of its essential role in DNA replication, ORC is required for every cell division during development. Thus, it is unclear how the Meier-Gorlin syndrome mutations in ORC lead to the tissue-specific defects associated with the disease. To address this question, we have used Cas9-mediated genome engineering to generate a *Drosophila melanogaster* model of individuals carrying a mutation in *ORC4*. Like the people with Meier-Gorlin syndrome, these flies reach adulthood, but have several tissue-specific defects. Genetic analysis revealed that this allele is a hypomorph and that mutant females are sterile. We demonstrated that this sterility is caused by a failure in DNA replication. By leveraging the well-studied Drosophila system, we showed that a disease-causing mutation in *orc4* disrupts DNA replication, and we propose that in individuals with MGS defects arise preferentially in tissues with a high-replication demand.

## Introduction

Meier-Gorlin syndrome (MGS) is a rare developmental disorder. Individuals with MGS show developmental defects, including primordial dwarfism, small or missing patella, and small ears (Bicknell *et al.* 2011b; Guernsey *et al.* 2011; De Munnik *et al.* 2012). A significant number of patients also present with microcephaly, though typically have normal cognitive function (De Munnik *et al.* 2012). The first case of MGS was reported in 1959 (Meier *et al.* 1959) with a second case following in 1975 (Gorlin *et al.* 1975), but the underlying genetic cause of the disease was unknown. Recently, advancements in next-generation sequencing have enabled the identification of mutations causing MGS. The identified mutations are in a set of genes (*ORC1, ORC4, ORC6, CDT1, CDC6*, *GMNN, CDC45*, and *MCM5*) responsible for the function of DNA replication origins, the chromosomal positions required for the initiation of DNA replication (Bicknell *et al.* 2011b; Guernsey *et al.* 2011; De Munnik *et al.* 2012; Burrage *et al.* 2015; Fenwick *et al.* 2016; Vetro *et al.* 2017).

The average human undergoes 10^16^ cell divisions in a lifetime, and every cell division requires faithful duplication of the genome. Genome duplication begins at multiple individual DNA replication origins that are formed in a cell-cycle regulated multistep process requiring many proteins that are conserved throughout eukaryotic organisms (Remus and Diffley 2009). In G1-phase, origins are selected by the binding of the origin recognition complex (ORC) comprised of six highly conserved members (Orc1, Orc2, Orc3, Orc4, Orc5, and Orc6). ORC recruits Cdc6 and together this complex recruits the Cdt1 chaperone bound to the MCM hexamer, the replicative helicase. In an ATP-dependent process, an MCM complex, comprised of two head-to-head hexamers (dhMCM), is formed on origin DNA, effectively ‘licensing’ the origin (Stillman 2005; Sclafani and Holzen 2007; Remus *et al.* 2009). In S-phase, multiple proteins, including S-phase kinases and the MCM helicase accessory factors, Cdc45 and GINS, convert the MCM complex into two active replicative helicases, which culminate in the initiation of DNA replication (origin function) (Moyer *et al.* 2006; Ilves *et al.* 2010). In most cell divisions, the genome must be replicated exactly once, and the cell-cycle separation of origin licensing (G1) and origin activation (S) ensures that only one complete round of genome duplication occurs per cell division (Deshaies 1995). However, in addition to this standard form of cell division, some cell-types undergo multiple rounds of genome duplication to generate polyploid cells (Lee *et al.* 2009). Both types of cell divisions depend on the same proteins for origin function.

As expected, based on the requirement for origin licensing for every cell division, null mutations in genes encoding proteins required for these processes are lethal (Micklem *et al.* 1993; Bell *et al.* 1993; Landis *et al.* 1997; Pinto *et al.* 1999; Pflumm and Botchan 2001; Shu *et al.* 2008; Park and Asano 2008; Baldinger and Gossen 2009; Balasov *et al.* 2009; Guernsey *et al.* 2011; Okano-Uchida *et al.* 2018). Thus, mutations in the genes that cause MGS must either be hypomorphic for their DNA replication functions or affect as yet undefined non-essential roles. Because origin function is essential in every cell division, it is unclear how MGS mutations that affect origin licensing result in tissue-specific defects. While ORC is essential for origin licensing, individual ORC subunits also function in other biological processes, such as heterochromatin formation (Prasanth *et al.* 2010) and cilia development (Hossain and Stillman 2012; Stiff *et al.* 2013). Thus, it is possible that defects in these other processes drive the MGS developmental disorder.

To determine the molecular mechanism underlying MGS mutations, MGS models have been generated in several organisms. In particular, the Orc4 MGS mutation has been generated in *Saccharomyces cerevisiae* (Sanchez *et al.* 2017). Yeast with this Orc4 (Y232C) substitution grow slowly due to defects in replication of the ribosomal DNA (rDNA) locus that normally contains hundreds of copies of the 9kb rDNA locus, each with its own origin. In *orc4*^*Y232C*^ yeast cells, the rDNA origin is insufficiently functional leading to DNA replication stress that causes substantial decreases in rDNA copy number, and the reduced translational capacity is not able to support a normal growth rate. Thus, while the *orc4*^*Y232C*^ allele clearly causes DNA replication defects at some yeast origins, it is challenging to determine whether the slow-growth phenotype is due to replication defects *per se* or the downstream effects on translation. In *Drosophila*, a transgenic system has been used to model MGS mutations in Orc6 (Bleichert *et al.* 2013; Balasov *et al.* 2015). These flies have several tissue-specific defects and biochemical analysis provides evidence that these phenotypes are due to a destabilization of ORC, which in turn results in decreased recruitment of the MCM hexamer to chromatin (Bleichert *et al.* 2013). Replication defects are also evident in cultured cells derived from MGS patients with multiple different *ORC1* alleles (Hossain and Stillman 2012). In contrast to the replication defects identified in yeast and flies, cell culture and zebrafish models for Orc1 mutants show defects in cilia development and these defects in turn may generate the various morphological phenotypes observed in transgenic fish models. (Bicknell *et al.* 2011b; Hossain and Stillman 2012; Yao *et al.* 2017; Maerz *et al.* 2019). Thus, while these models have provided important insights into this pleotropic disease, it remains unclear whether the different mechanisms identified reflect differences in the underlying causes of MGS or whether they reflect differences in how the mutations were modelled. Finally, it is also important to note that the metazoan models to date rely on exogenous expression of the mutant protein and therefore do not precisely mimic the conditions observed in MGS individuals.

To better understand the molecular mechanisms underlying the tissue-specific phenotypes caused by MGS mutations, we used Cas9-genome editing to generate a *Drosophila* model of MGS. Because there is a single identified MGS mutation in a highly conserved region of Orc4 (Y174C), we generated a *Drosophila* model for this mutation and demonstrated that similar to the disease phenotype, flies homozygous for this mutation are viable. In addition, we created a wild-type control and a null mutant allele, which enabled us to demonstrate that the MGS mutation is a hypomorphic allele that causes tissue-specific replication defects resulting in female sterility. Together our data suggest that the tissue-specific defects identified in MGS patients may result from cells within these tissues having a high-replication demand that cannot be met by the MGS mutant replication factors. Further, this work demonstrates the power of genome-editing in the fly to model human disease.

## Materials and Methods

### Fly lines and husbandry

Flies were grown on standard molasses food at 25°C. Fly lines used in this study: *orc4*^*WT*^ (this study), *orc4*^*Y162C*^ (this study), *orc4*^*null*^ (this study), *mcm6*^*K1214*^ *v[24]/FM3* (Bloomington *Drosophila* Stock Center (BDSC) #4322), *hsFlp122;Sp/SM6-TM6b*, *FRTG13 orc4*^*Y162C*^*/CyO* (this study), and *P{w[+mW.hs]=FRT(w[hs])}G13 P{w[+mC]=ovo*^*D1-18*^*}2R/T(1;2)OR64/CyO* (BDSC #4434).

### Generation of *orc4*^*WT*^ and *orc4*^*Y162C*^ fly lines

Single-stranded donor oligonucleotides (ssODNs) were generated to target the region of *orc4* encoding Y162. Each ssODN had silent mutations to mutate the PAM site and generate a novel *Nde*I cut site for molecular screening. The *orc4*^*Y162C*^ ssODN also contained the necessary alterations to create the Y162C mutation. The *orc4*^*WT*^ ssODN did not contain this mutation. gRNA plasmids and ssODNs were mixed, injected, and screened as in (Hamm *et al.* 2017). Injections were done by Best Gene Inc.

*orc4*^*WT*^ ssODN:

AAGAGAAAGACCTGCCGGTGCGAGAAACGCGACTTGACCCGCTTCTCCAGCAGCTCGATCACGTCGAGGCGACAGGTAACGCCAAGTACACATATGGGCGCCTGGGCTACTGGGAGACGTCGAAGAGGTTGTAAAGCAGGGTCTGGTTGTGGTGAGCACAGAAGAGGTCGAACTCCTCGAGAAT

*orc4*^*Y162C*^ ssODN:

AAGAGAAAGACCTGCCGGTGCGAGAAACGCGACTTGACCCGCTTCTCCAGCAGCTCGATCACGTCGAGGCGACAGGTAACGCCAAGTACACATATGGGCGCCTGGGCTACTGGGAGACGTCGAAGAGGTTGCAAAGCAGGGTCTGGTTGTGGTGAGCACAGAAGAGGTCGAACTCCTCGAGAAT

Forward Screening Primer: GAAGTCCATCACTGTGCAGAT

Reverse Screening Primer: TGGTTGCGGGAGAAGTAAAG

### Viability Assays

3-5 heterozygous males and 5-10 heterozygous females of the indicated genotypes were mated in standard molasses vials with dry yeast and flipped twice at 2-day intervals. Two days after the final flip, the adult flies were cleared from the vials and their progeny were allowed to reach adulthood. Over 800 adults were counted for each cross. The ratio of CyO and non-CyO adults was determined and the *χ*^2^ value was calculated for each cross, correcting for the observed ratio from the *orc4*^*WT*^/*CyO* self-cross.

### Adult Phenotyping

Adult flies from the indicated genotypes were imaged on a Nikon SMZ745 dissection microscope (back bristles) or frozen at −20°C and then imaged on a Zeiss Axioplan2 epifluorescence microscope (wing bristles).

### Ovary DAPI Staining

Females of the indicated genotypes were mixed with males in molasses vials with a small amount of yeast paste and grown for two days. Flies were flipped into a fresh vial after 24 hours. Ovaries were dissected into Grace’s medium. The media was removed, and the ovaries were resuspended in 0.5 ml of fix solution (4% formaldehyde in 1xPBS) and incubated for 15 minutes at room temperature while rocking. The ovaries were washed twice with 1ml of PBST (1XPBS + 0.2% Triton X-100) and then washed for 5 minutes with 1XPBS to remove the detergent. Ovaries were then incubated with 1xPBS + DAPI (1:1000) for 15 minutes and then mounted on a slide, covered with a coverslip, and sealed with clear nail polish. Ovaries were imaged on Zeiss Axioplan2 epifluorescence microscope.

### Nurse Cell Counts

DAPI stained ovaries from the indicated genotypes were imaged on a Zeiss Axioplan2 epifluorescence microscope, and the nurse cells were counted. Nurse cells from 50-100 stage 10 egg chambers were counted for each genotype.

### EdU Assay

Ovaries from ten females were dissected as described above and resuspended in 100 ul of Grace’s media. 100 ul of 2XEdU in Grace’s media (15 uM) stock solution was added to each sample and incubated for 1.25 hours at room temperature. The ovaries were washed twice with 200 ul of 3% BSA in 1XPBS for 5 minutes each time. The ovaries were then fixed for 15 minutes in 200 ul of 4% formaldehyde in 1XPBS. After fixation, the ovaries were washed twice with 200 ul of PBST (1XPBS + 0.5% Triton X-100) for 5 minutes and 20 minutes. Ovaries were then washed twice for 5 minutes each with 1XPBS + 3% BSA and then carried through the Click-iT Plus EdU Imaging Kit protocol (Thermo Fisher Scientific). Ovaries were imaged on a Nikon A1R-SI+ confocal microscope and >50 stage 10 chambers were examined for EdU foci in each genotype.

### Generation of Germline Mitotic Clones and Egg Counting

*FRTG13 orc4*^*Y162C*^/*CyO* females were mated with *hsFLP122; P{w[+mW.hs]=FRT(w[hs])}G13 P{w[+mC]=ovoD1-18}2R/T(1;2)OR64/CyO* males. Their progeny were incubated for 30 minutes at 37°C in a circulating water bath either 24-48 hours after egg laying (1x heat shock) or 24-48 and 48-72 hours after egg laying (2x heat shock). These embryos were reared to adulthood. Non-CyO females were isolated and mated to males in standard molasses vials. Homozygous *orc4*^*WT*^ and *orc4*^*Y162C*^ females were also mated to males as controls. The crosses were flipped every 24 hours for four days, and the number of eggs laid each day were counted.

### Data availability

Strains and plasmids are available upon request. The authors affirm that all data necessary for confirming the conclusions of the article are present within the article, figures and tables.

## Results

### Generating endogenous point mutations in *Drosophila* to model Meier-Gorlin syndrome

Meier-Gorlin syndrome (MGS) results from mutations in multiple different origin regulatory proteins, each of which is required for replication during every cell division of the organism (Bicknell *et al.* 2011b; Guernsey *et al.* 2011; De Munnik *et al.* 2012). Given the additional cellular roles of these factors, we sought to model the disease in a rapidly developing metazoan to better understand the mechanisms behind the phenotypic defects. The only mutation in *ORC4* that causes MGS is the substitution of tyrosine 174 to cysteine (Guernsey *et al.* 2011). Tyrosine 174 is in the AAA+ ATPase domain of ORC4 (Figure 1A). This region is highly conserved from yeast to humans, suggesting that it plays a critical role in ORC function and enabled us to identify the homologous residue (tyrosine 162) in *Drosophila*.

**Figure 1:**
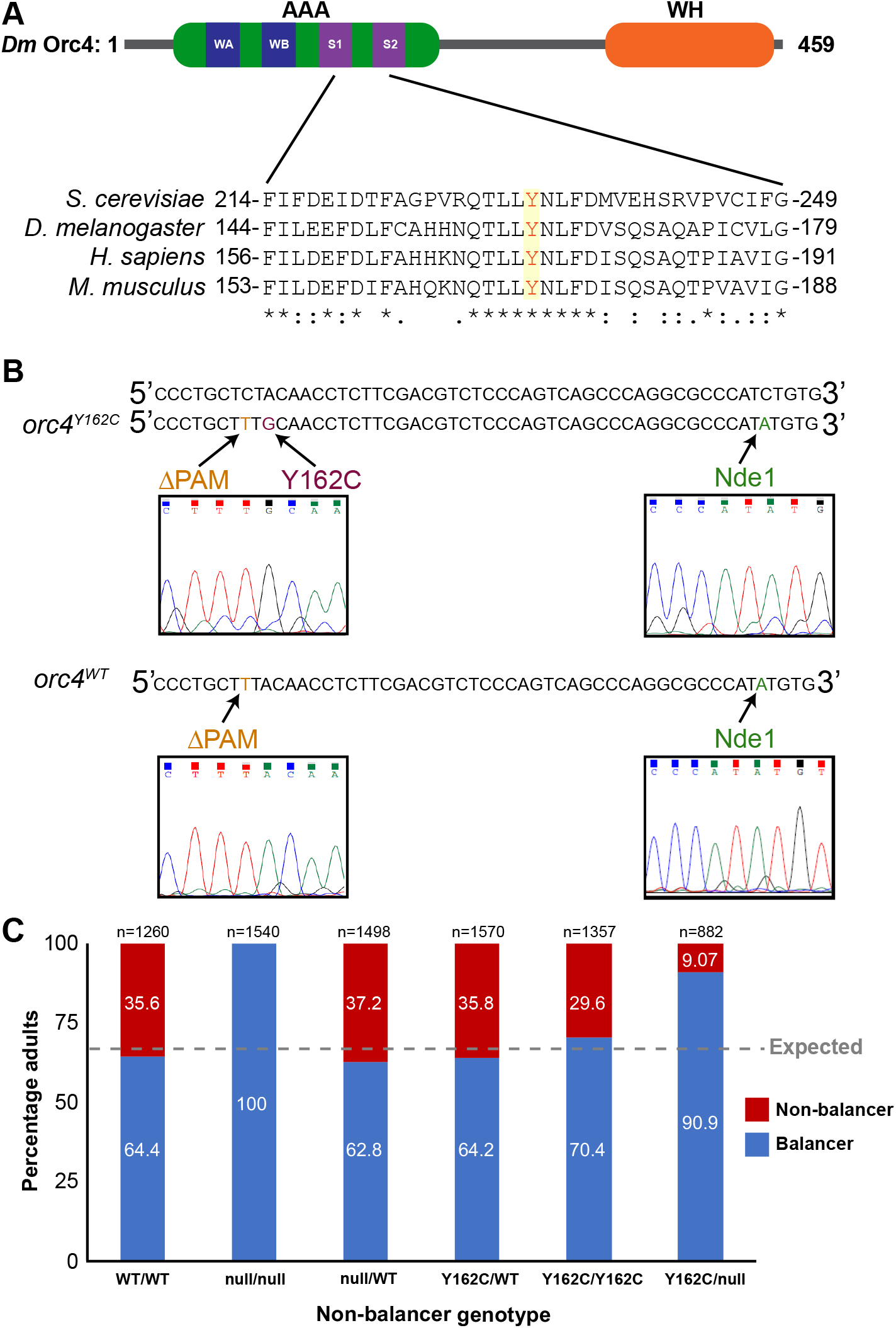
Similar to MGS individuals, *orc4*^*Y162C*^ animals are viable. (A) Schematic of *Drosophila* Orc4 protein domains with the AAA+ ATPase domain (AAA) and the winged helix (WH) domain indicated. The Walker A (WA), Walker B (WB), Sensor 1 (S1) and Sensor 2 (S2) motifs are also indicated in the AAA+ ATPase domain (*top*). Clustal Omega alignment of multiple eukaryotes for the region surrounding the conserved tyrosine residue mutated in MGS (highlighted). (*bottom*) (B) Sequence of the genomic region encoding Y162 of Orc4 from a wild-type, unedited *Drosophila* strain (top). Below the sequence of the Cas9-edited genomes of the *orc*^*Y162C*^ and *orc*^*WT*^ strains are shown. The silent mutation removing the PAM site (gold), the Y162C mutation (maroon), and silent *Nde*I cut site (green) are noted along with the sequencing traces confirming the edited alleles. (C) The percent of balancer (CyO) to non-balancer adults are shown for the crosses resulting in the indicated non-balancer progeny. Heterozygote males and females were mated, and their progeny were scored for the presence of the CyO balancer. The dashed gray line represents the expected ratio from this cross (66% balancer: 33% non-balancer). n = total number of flies assayed.

We used Cas9-mediated genome engineering to introduce a mutation encoding a tyrosine to cysteine substitution (Y162C) at the endogenous *orc4* locus. Mutating the endogenous locus assured that any phenotypic changes observed were due to the mutation and not to changes in transcription that might be caused by transgenic constructs. In addition to the Y162C substitution, our genome engineering strategy introduced a silent mutation in the protospacer adjacent motif (PAM) to inhibit the guide RNA from directing cleavage of the engineered genome. We also engineered a silent mutation introducing a *Nde*I restriction site to allow for rapid molecular screening of the modified genomes (Figure 1B). To ensure that the two silent mutations do not affect Orc4 activity, we also generated a control strain, *orc4*^*WT*^, which does not have the Y162C substitution, but includes both silent mutations. In addition, our editing generated a likely null allele, *orc4*^*null*^, through a single base pair deletion resulting in a frameshift after I187. While we have not analyzed protein product from this allele, we presume that this is a null allele based on the molecular nature of the mutation and the fact that, similar to null alleles in other ORC subunits, homozygous *orc4*^*null*^ animals are not viable (Landis *et al.* 1997; Pinto *et al.* 1999; Pflumm and Botchan 2001; Park and Asano 2008; Baldinger and Gossen 2009; Balasov *et al.* 2009). Together, these strains provide the first metazoan model for an MGS mutation in which the mutation was engineered at the endogenous locus along with precisely defined control strains.

### orc4^Y162C^ is a hypomorph and animals are homozygous viable

MGS individuals with *ORC4*^*Y174C*^ mutations are either homozygous for this mutation or carry it over a null allele (Bicknell *et al.* 2011a; Guernsey *et al.* 2011; De Munnik *et al.* 2012). While these individuals possess a number of distinctive phenotypes, they survive and have a normal expected lifespan. Therefore, we initially tested the viability of homozygous *orc4*^*Y162C*^ animals. We quantitatively assessed the number of non-balancer (straight-winged) and balancer (curly winged) progeny produced from crosses of balanced heterozygotes for either *orc4*^*Y162C*^, *orc4*^*WT*^ or *orc4*^*null*^ heterozygotes. Because animals homozygous for the CyO balancer die as larvae, Mendelian ratios would predict that 66% of the adults would carry the balancer while the remaining 33% would not. As expected, a single wild-type copy of *orc4* resulted in ratios close to those expected (Figure 1C). Furthermore, *orc4*^*null*^ animals are inviable, similar to previously reported data for null alleles of other ORC members (Landis *et al.* 1997; Pinto *et al.* 1999; Pflumm and Botchan 2001; Park and Asano 2008; Baldinger and Gossen 2009; Balasov *et al.* 2009) (Figure 1C). Like individuals carrying the *ORC4*^*Y174C*^ mutation, *orc4*^*Y162C*^ animals are homozygous viable. Unlike some prior models of MGS mutations (Bicknell *et al.* 2011b; Yao *et al.* 2017; Maerz *et al.* 2019), we did not observe any reduction in size of the homozygous *orc4*^*Y162C*^ animals, suggesting either a difference between *Drosophila* and other organisms or that some of these phenotypes may have been caused by misexpression of the disease allele which was avoided by our genome-editing strategy. When we corrected our expected ratio of *CyO* to non-*CyO* flies based on the observed ratio from the *orc4*^*WT*^/*CyO* cross, we identified a statistically significant decrease in the number of *orc4*^*Y162C*^/*orc4*^*Y162C*^ adults (χ^2^, p=5.0×10^−6^).

The observed impact on viability of the homozygous *orc4*^*Y162C*^ mutation suggested it might be a loss-of-function allele. To directly test this, we scored the viability of *orc4*^*Y162C*^/ *orc4*^*null*^ trans-heterozygotes. These flies reach adulthood at reduced levels as compared to *orc4*^*Y162C*^; we recovered one third of the expected number of trans-heterozygotes (χ^2^, p=1.1×10^−60^), demonstrating that *orc4*^*Y162C*^ is a hypomorphic allele. Together, these data showed that similar to MGS patients, the homozygous *orc4*^*Y162C*^ animals are viable. Furthermore, our genetic analysis demonstrated that the tyrosine to cysteine substitution in Orc4 likely results in a protein with reduced functionality.

### Animals homozygous for *orc4*^*Y162C*^ have tissue-specific defects

Given that ORC is required during replication of every cell, an outstanding question regarding the phenotypes of individuals with MGS is why there are tissue-specific defects. Having demonstrated that *orc4*^*Y162C*^ animals reach adulthood, we were able to assay for tissue-specific phenotypes. When compared to a wild-type strain (*w^1118^*), *orc4*^*WT*^ animals show no obvious phenotypic differences. Thus, we used these as our wild-type controls throughout the remainder of the study. By comparison, *orc4*^*Y162C*^ homozygous animals had several phenotypic abnormalities. We identified several missing bristles on the thorax (Figure 2A). In addition, we observed severe bristle defects on the wing. The bristles along the wing are normally uniform in length and evenly spaced. By contrast, the wing margin bristles in the *orc4*^*Y162C*^ animals are disorganized and vary in length along the wing (Figure 2A). Therefore, *orc4*^*Y162C*^ homozygous animals resemble individuals with MGS as they possess tissue-specific defects, further strengthening the relevance of our *Drosophila* model.

**Figure 2:**
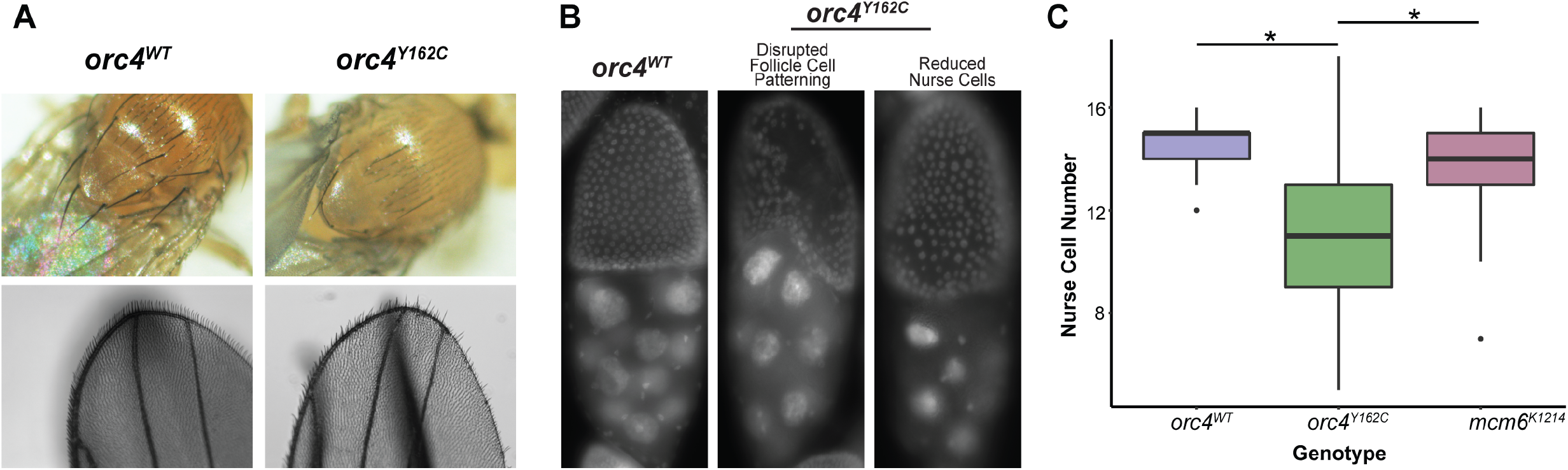
*orc4*^*Y162C*^ animals have tissue-specific phenotypes. (A) Images of homozygous *orc4*^*WT*^ and *orc4*^*Y162C*^ animals. Mutant animals have missing back bristles. (*top*) and absent as well as disorganized wing-margin bristles (*bottom*). B. DAPI stained images of stage 10 egg chambers from homozygous *orc4*^*WT*^ and *orc4*^*Y162C*^ animals. Examples of *orc4*^*Y162C*^ animals with disorganized follicle cells and reduced nurse cells are shown. (C) Quantification of the number of nurse cells in *orc4*^*WT*^, *orc4*^*Y162C*^, and *mcm6*^*K1214*^ ovaries. *orc4*^*Y162C*^ females have significantly fewer nurse cells than both *orc*^*WT*^ (p < 2.7 x10^−17^ *t*-test) and *mcm6*^*K1214*^ females (p<1.0 x10^−10^, *t*-test). Fifty stage 10 chambers were counted in *orc4*^*WT*^ and *mcm6*^*K1214*^ animals, 100 stage 10 chambers were counted in *orc4*^*Y162C*^ animals.

### Females homozygous for orc4^Y162C^ are sterile

While *orc4*^*Y162C*^ animals were homozygous viable, females were sterile. By contrast, homozygous *orc4*^*Y162C*^ males were fertile. Females produced eggs at very low frequencies compared to *orc4*^*WT*^ control animals. The few eggs produced did not have dorsal appendages and appeared watery and malformed, indicative of a thin eggshell. Indeed, female sterility is a phenotype shared amongst animals possessing loss-of-function mutations in a variety of replication factors, such as Orc2, MCM6, Cdt1, and chiffon (Orr-Weaver 1991; Landis *et al.* 1997; Landis and Tower 1999; Whittaker *et al.* 2000).

Because the females are sterile, we examined the ovaries of *orc4*^*Y162C*^ homozygous animals to determine if there were specific defects in egg chamber development. We dissected ovaries from *orc4*^*WT*^ and *orc4*^*Y162C*^ homozygous females and stained them with 4’, 6-diamidino-2-phenylindole (DAPI) to image the nuclei. The ovarioles appeared largely normal, and we could identify egg chambers through stage 14. Nonetheless, we noted two distinct phenotypes in stage 10 chambers: disrupted follicle-cell patterning and decreased numbers of nurse cells (Figure 2B). Both cell types play critical roles in oocyte maturation, and the deleterious phenotypes we observed could be responsible for the female sterility. The somatically derived follicle cells rapidly amplify the chorion genes during stage 10B (Orr-Weaver 1991). Simultaneously, the nurse cells produce the maternally derived products that will be deposited into the egg. Both cell types are polyploid, and the disorganized follicle cell structure and decreased nurse cell numbers could potentially be due to replication defects during oocyte maturation.

To more quantitatively assess these defects, we determined the number of nurse cells in *orc4*^*WT*^, *orc4*^*Y162C*^, and *mcm6*^*K1214*^ homozygous females. We included *mcm6*^*K1214*^ as a control since it is a female-sterile allele of an additional component of the replication machinery (Komitopoulou *et al.* 1983). Using fixed, DAPI stained ovaries, we counted the number of nurse cells in 50-100 stage 10 egg chambers. Wild-type stage 10 egg chambers possess 15 nurse cells. Ovaries from *orc4*^*Y162C*^ homozygous females have fewer nurse cells than *orc4*^*WT*^ females along with a wider distribution in the number of nurse cell per egg chamber (*t*-test, p=4.16×10^−12^) (Figure 2C). By contrast, stage 10 chambers from the *mcm6^K1214^* females showed only a minor decrease in nurse cell number as compared to ovaries from *orc4*^*WT*^ females *(t-*test, p=7.79×10^−3^) and were significantly different in comparison to *orc4*^*Y162C*^ (*t*-test, p=1.03×10^−10^). Thus, while both the *orc4*^*Y162C*^ and *mcm6*^*K1214*^ alleles lead to female sterility, only *orc4*^*Y162C*^ females have a decreased number of nurse cells. This result provides evidence that the sterility of *orc4*^*Y162C*^ females may result from a defect in cells distinct from the nurse cells.

### Females homozygous for *orc4*^*Y162C*^ fail to amplify the chorion genes

Mutations in genes encoding replication factors are known to cause female sterility at least in part due to replication defects in the somatic follicle cells. At stage 10B, follicle cells in the egg chamber undergo endoreplication, selectively amplifying a limited subset of loci including the chorion genes, which are essential for eggshell production later during oocyte maturation (Orr-Weaver 1991; Calvi *et al.* 1998). This results in a gene amplification of 16-20 fold for a region on the X chromosome and 60-80 fold for a region of the 3^rd^ chromosome. Amplification occurs through repeated rapid and precise rounds of origin firing and replication fork elongation. Failure to adequately amplify these loci leads to thin, fragile eggshells, which results in female sterility. Thus, the female sterility of *orc4*^*Y162C*^ may result from a failure to adequately amplify the chorion genes during oocyte maturation. Furthermore, well-established assays for chorion gene amplification provide a system by which to directly assay whether the MGS mutation in *orc4* impacts DNA replication (Park and Asano 2012).

To test if *orc4*^*Y162C*^ animals are replication deficient, we dissected ovaries from *orc4*^*WT*^, *orc4*^*Y162C*^, and *mcm6*^*K1214*^ homozygous females and incubated them with the modified thymidine analogue EdU for 1.25 hours, which allowed for the incorporation of EdU into replicating DNA that could subsequently be imaged using a small molecule-based fluorescent assay. Amplification of the chorion gene loci can be visualized as distinct EdU foci in the follicle cells of stage 10B egg chambers. Indeed, in the *orc4*^*WT*^ females we observed large robust foci in the follicle cells (Figure 3A). As expected, no foci were detected in stage 10 egg chambers from *mcm6*^*K1214*^ ovaries (Figure 3A). Despite clear incorporation of EdU in other stages of egg chamber development, no EdU foci were evident in the stage 10 egg chambers of *orc4*^*Y162C*^ females (Figure 3A). To quantify this replication defect, we imaged >50 stage 10 egg chambers for each genotype. There were large robust foci in 45% of the *orc4*^*WT*^ stage 10 egg chambers, but none in *orc4*^*Y162C*^ animals (*t*-test, p=2.67×10^−11^ vs. *orc4*^*WT*^) (Figure 3B). In ovaries from *mcm6*^*K1214*^ females, we identified weak, fractured foci in 2% of the stage 10 egg chambers (*t*-test, p=4.60×10^−8^ vs. *orc4*^*WT*^). We noted that the Hoechst signal for the *orc4*^*Y162C*^ animals was lower as compared to either the *orc4*^*WT*^ or *mcm6*^*K1214*^ animals. While this could be an issue with the staining, all the samples were stained at the same time with the same reagent mix. Thus, it is possible that this reflects a lower DNA content in the follicle cells of the *orc4*^*Y162C*^ females as compared to control animals. Because the follicle cells themselves undergo a few rounds of total endoreplication cycles, it is possible this difference in Hoechst staining reflects a decrease in this endoreplication. Together these data provide evidence that the *orc4*^*Y162C*^ mutation leads to a specific replication defect as females fail to amplify the chorion gene locus and this in turn results in the observed thin, fragile eggshells. Thus, while *orc4*^*Y162C*^ mutant animals can replicate their DNA, DNA replication is defective during chorion gene amplification where there may be a high demand for rapid origin licensing and firing.

**Figure 3:**
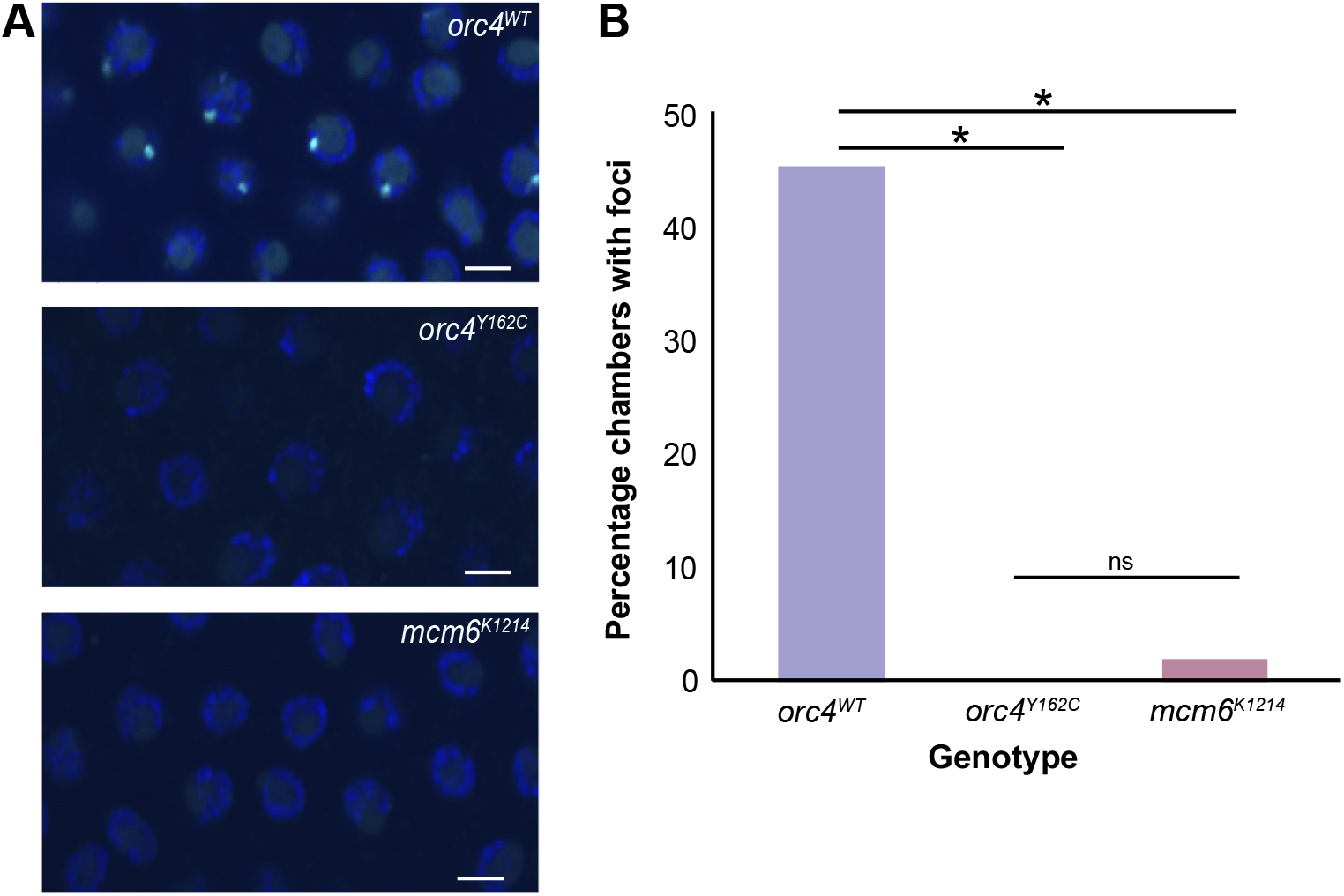
*orc4*^*Y162C*^ females fail to replicate the chorion gene loci. (A) EdU staining of stage 10B egg chambers from *orc4*^*WT*^, *orc4*^*Y162C*^, and *mcm6*^*K1214*^ females. EdU staining (cyan) marks the amplified chorion genes and was acquired with identical settings for all genotypes. DNA is stained with Hoechst (blue). Scale bar, 5 µm. (B) Quantification of stage 10 chambers with EdU foci from *orc4*^*WT*^, *orc4*^*Y162C*^, and *mcm6*^*K1214*^ females. *orc4*^*Y162C*^ and *mcm6*^*K1214*^ animals fail to amplify the chorion genes as compared to *orc4*^*WT*^ females (p<1×10^−5^, *t*-test). More than 50 stage 10 chambers were assayed for each genotype.

### Animals inheriting maternal *orc4*^*Y162C*^ cannot complete embryogenesis

Apart from endoreplication, another distinct developmental time point that requires rapid origin licensing and firing is during the synchronous nuclear divisions in the early embryo. Immediately following fertilization, development is controlled by maternally deposited products while the genome is reprogrammed. During this time, the nuclei are in a shared, syncytial cytoplasm and divide quickly with an abbreviated cycle comprised of only a synthesis (S) phase and mitosis. These divisions occur approximately every 10 minutes with DNA being replicated in about half of this time. We hypothesized that if the tyrosine-to-cysteine mutation in Orc4 caused defects in tissues in which there was a high demand for DNA replication that embryos inheriting maternal *orc4*^*Y162C*^ would fail to progress through the early stages of development. To address this, we used the FLP/FRT system to generate germline clones that are homozygous for the *orc4*^*Y162C*^ mutation, while remaining largely heterozygous for the mutation in the somatic follicle cells (Chou and Perrimon 1996). Because we combined this with the dominant female-sterile *ovo*^*D1*^ mutation (Chou *et al.* 1993), this strategy generated animals that only inherited *orc4*^*Y162C*^ maternally. Females heterozygous for both the *ovo*^*D1*^ and the *orc4*^*Y162C*^ mutations did not produce eggs, as expected. By contrast, heat-shocked females laid some eggs, and these eggs had dorsal appendages, a striking difference from the very few eggs laid by *orc4*^*Y162C*^ homozygous females (Figure 4A). This suggests that our strategy at least partially rescued eggshell production. We quantified the numbers of eggs laid by females in which the germline clones (glc) were generated. These females laid ∼4 times more eggs than *orc4*^*Y162C*^ females (Figure 4B). While these embryos were rescued for eggshell production, they failed to complete embryogenesis and showed general morphological defects (Figure 4A). This failure to progress through embryogenesis further supports the model that the hypomorphic *orc4*^*Y162C*^ mutation results in replication defects in tissues that require rapid origin licensing and firing and that it is this tissue-specific requirement that leads to the distinct phenotypes of MGS individuals.

**Figure 4:**
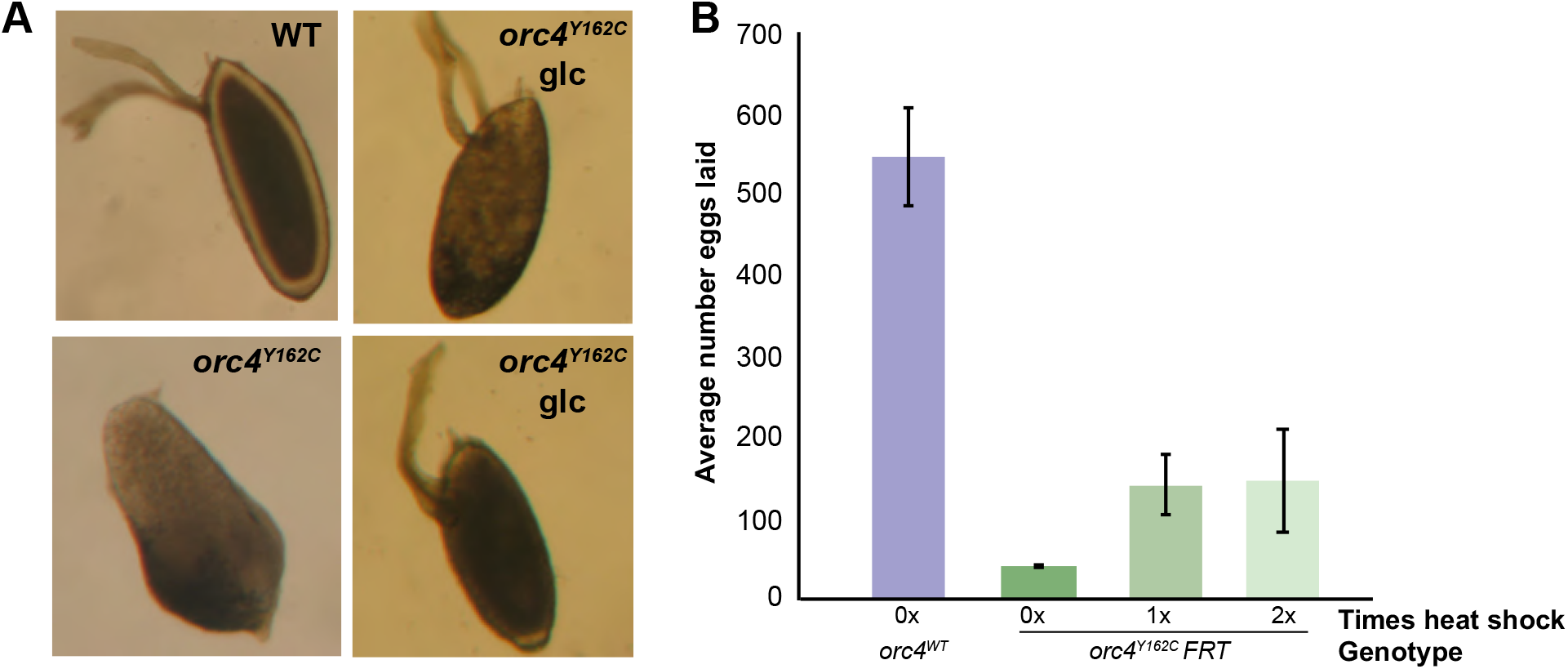
Embryos inheriting maternal orc4^Y162C^ are inviable. (A) Images of embryos laid by mothers of the indicated genotype. WT (wild-type females); *orc4*^*Y162C*^ (*orc4*^*Y162C*^ females); *orc4*^*Y162C*^ *glc* (heat shocked *hsFLP122; FRTG13 ovo*^*D1*^/*FRTG13 orc4*^*Y162C*^ females with germline clones). Dorsal appendages are evident in embryos laid by wild-type females and females heterozygous for *orc4*^*Y162C*^ in the follicle cells (*orc4*^*Y162C*^ *glc*), but not from females homozygous for *orc4*^*Y162C*^. (B) Average numbers of eggs laid from ten females of the indicated genotypes over four days are shown. Error bars indicate the standard deviation from three biological replicates.

## Discussion

The underlying mechanisms generating the phenotypes of MGS individuals are not well understood. While the mutations that lead to this disease are clustered in genes required for the licensing of origins of replication during the G1 phase of the cell cycle, ORC has additional functions outside of DNA replication. Using Cas9-mediated mutagenesis, we have engineered the endogenous *orc4* locus to establish a metazoan model for MGS. Our *Drosophila* model recapitulates the disease state as animals are homozygous viable, demonstrating that the encoded tyrosine-to-cysteine mutation in Orc4 is compatible with DNA replication in many tissues. Genetic analysis showed that *orc4*^*Y162C*^ is a hypomorph that is predicted to generate a protein with reduced functionality. Indeed, *orc4*^*Y162C*^ females are sterile, and this is likely due to a replication defect in tissues with a high-replication demand, like the ovarian follicle cells. Together our data provide a mechanistic model for MGS phenotypes and, in so doing, demonstrate the utility of the well-established *Drosophila* system for rapidly generating metazoan disease models and testing specific mechanistic hypotheses to explain the observed patient phenotypes.

In this study, we demonstrate that the *orc4*^*Y162C*^ mutation results in defects in two distinct tissues with high-replication demand (the early embryo and chorion gene amplification in the follicle cells). Based on these data, we propose that the MGS mutation in Orc4 generates a sub-functional ORC that cannot meet the replication demands, resulting in defects specifically in tissues requiring fast/efficient replication, which are particularly sensitive to compromised replication machinery. While we propose that replication speed is a critical factor that leads to the phenotypic defects in *orc4*^*Y162C*^ animals, we cannot exclude the possibility that the defects are due to the non-canonical cell cycles in the tissues assayed. In the nurse and follicle cells of the ovary and in the early embryo, the replication cycle does not occur with the standard, four phases (G1, S, G2, M). Instead, the nurse cells undergo multiple rounds of complete genome endoreplication. In the follicle cells, the chorion gene loci undergo multiple rapid rounds of re-initiation of DNA synthesis. Similarly, in the early embryo the nuclear division cycle is a series of rapid synthesis and mitosis phases without gap phases. Further experiments will be required to determine if phenotypes caused by MGS mutations are due specifically to defects in tissues with cycles of rapid DNA replication, non-standard cycles of DNA replication, or both.

What is clear is that *orc4*^*Y162C*^ flies are defective in replicating the chorion gene loci and that this causes female sterility. Similarly, yeast with the corresponding tyrosine-to-cysteine mutation (Y232C) show replication defects, particularly at the ribosomal DNA locus origin (Guernsey *et al.* 2011; Sanchez *et al.* 2017). Only this tyrosine-to-cysteine mutation at position 174 has been associated with MGS, suggesting that a very limited subset of mutations can be tolerated for viability and that this may be the one of few permissible mutations at this locus. From recent structural studies, it is evident that the region of Orc4 surrounding and including Y174 interacts with the ATPase domain of Orc1 with Y174 of Orc4 making contacts with E621 of Orc1 (Tocilj *et al.* 2017). This domain of Orc4 is highly conserved across species, suggesting there is significant evolutionary constraint placed on these residues. This likely reflects a functional requirement for the interaction between Orc4 and Orc1 in this region. Indeed, the tyrosine-to-cysteine mutation in human Orc4 results in altered ORC ATPase activity *in vitro* (Tocilj *et al.* 2017). Because ORC ATPase activity is essential for origin function, these data, combined with our *in vivo* observations, provide a compelling molecular model in which the *orc4*^*Y162C*^ mutation leads to a functional, but compromised ORC that results in decreased recruitment of the MCM hexamer to chromatin. Because most cells license more origins of replication than are necessary to replicate the genome, this reduction in MCM loading can be tolerated in most tissues. However, in tissues that require rapid or efficient rounds of replication, the decrease in MCM loading may result in replication defects. Indeed, limiting amounts of the MCM helicase can have tissue-specific defects as demonstrated by the fact that mice with a hypomorphic allele of *MCM3* die from anemia due to replication failure of the rapidly dividing erythrocyte precursor cells (Alvarez *et al.* 2015).

Together these data from multiple model systems suggest that in individuals with the Orc4 Y174C MGS mutation have an insufficient number of origins to support rapidly replicating tissues. Nonetheless, it remains unclear if MGS mutations in other ORC subunits similarly result in decreased replication capacity in specific tissues. Biochemical and phenotypic data from *Drosophila* suggest that the MGS mutation in *orc6* results in a replication defect caused by a destabilization of ORC (Bleichert *et al.* 2013; Balasov *et al.* 2015). Similar to the model we propose for Orc4 Y174C, this destabilized ORC decreases the recruitment of the MCM hexamer to chromatin (Bleichert *et al.* 2013). Modelling of the *Orc1* MGS mutations in zebrafish generates small fish with morphological defects, similar to the phenotypes in MGS individuals. In contrast to the models proposed for Orc4 and Orc6, the phenotypes in the zebrafish model of *Orc1* MGS mutations have been suggested to arise from defects in cilia formation (Bicknell *et al.* 2011b; Yao *et al.* 2017; Maerz *et al.* 2019). Thus, it remains possible that roles for ORC proteins outside of origin licensing may impact MGS phenotypes. Future work in multiple organisms will be needed to identify how each MGS mutation leads to the disease phenotypes. Nonetheless, our data clearly demonstrate tissue-specific replication defects caused by the MGS-associated mutation in Orc4. Thus, the tissue-specific defects in MGS patients may arise because cells within these tissues have specialized replication demands. Furthermore, our data demonstrate the ability to gain mechanistic insights into disease phenotypes by combining rapid and precise editing of the *Drosophila* genome with the wealth of tools and knowledge derived from over a century of studying this powerful model metazoan.

